# Early changes in microRNA expression in Arabidopsis plants infected with the fungal pathogen *Fusarium graminearum*

**DOI:** 10.1101/2024.05.29.596347

**Authors:** Savio S. Ferreira, Suman Pandey, Jesseca Hemminger, Serdar Bozdag, Mauricio S. Antunes

## Abstract

Plants respond to biotic stressors by modulating various processes in an attempt to limit the attack by a pathogen or herbivore. Triggering these different defense processes requires orchestration of a network of proteins and RNA molecules that includes microRNAs (miRNAs). These short RNA molecules (20-22 nucleotides) have been shown to be important players in the early responses of plants to stresses because they can rapidly regulate the expression levels of a network of downstream genes. The ascomycete *Fusarium graminearum* is an important fungal pathogen that causes significant losses in cereal crops worldwide. Using the well-characterized *Fusarium-Arabidopsis* pathosystem, we investigated how plants change expression of their miRNAs globally during the early stages of infection by *F. graminearum*. In addition to miRNAs that have been previously implicated in stress responses, we have also identified evolutionarily young miRNAs whose levels change significantly in response to fungal infection. Some of these young miRNAs have homologs present in cereals. Thus, manipulating expression of these miRNAs may provide a unique path toward development of plants with increased resistance to fungal pathogens.

## Introduction

Because they cannot run or hide from threats, plants have developed exquisite ways to constantly survey their environment and respond accordingly to different stress conditions. These responses typically involve many layers, from activation of membrane-bound receptor proteins to up- or down-regulation of expression of genes that ultimately lead to a physiological change to cope with the stress. One such layer consists of different types of non-coding RNA molecules, which can control expression of a network of genes involved in the response to a particular stress.

Small RNAs (sRNAs), such as microRNAs (miRNAs) and small interfering RNAs (siRNAs), are 20-30 nucleotide-long non-coding RNA molecules that are involved in the sequence-specific regulation of gene expression at the transcriptional or post-transcriptional level. In addition to their roles in growth, development and maintenance of genome integrity, sRNAs are also important components in plant stress responses. Changes in the levels of endogenous plant sRNAs in response to external stimuli, such as pathogen infection, environmental conditions, and nutrient availability, are well-documented ^1–3^. These sRNA regulatory networks in plants are thought to have evolved as a cellular defense mechanism against RNA viruses and transposable elements that were later adapted to regulate the expression of endogenous genes ^4^. Abundant evidence now suggests that endogenous plant sRNAs and their activity in silencing target genes may serve as a general regulatory mechanism in plant immune responses to many pathogens ^5–7^. In fact, to try to counteract these responses, some bacterial effector proteins and viral proteins have been shown to suppress host miRNA biogenesis and/or activity ^8–10^. Therefore, several studies have looked at changes in the levels of plant sRNAs, and how they affect the protein-coding transcriptome, to identify common mechanisms involved in responses of plants to pathogens and pests ^11, 12^.

In plants, several miRNAs have been shown to be up-regulated, whereas others are down-regulated, during their interaction with pathogens and pests ^13–16^. Many of these miRNAs are predicted to function as early regulators of stress-induced transcription factors that, in turn, influence the expression of defense genes ^17^. For example, the levels of specific miRNAs have been shown to change during progression of the Huanglongbing (HLB) disease in citrus ^18^, infection of Arabidopsis plants by the bacterial pathogen *Pseudomonas syringae* ^19^, and the infection of Lilium plants by the fungal pathogen *Botrytis elliptica* ^13^. Furthermore, differential regulation of miRNAs in plants has been observed depending on whether they interact with a pathogen or beneficial organism, with differences also observed between roots and leaves ^14^.

Previous recent studies on the interactions between biotic factors and Arabidopsis plants have mostly focused on changes in protein-coding genes by RNA-Seq analysis ^20–24^. Changes in sRNAs are usually not considered, although a bioinformatics approach has been applied to identify miRNA promoters in Arabidopsis that are putatively bound and regulated by *Xanthomonas campestris* effector proteins ^25^. Although this study identified miRNAs that may be up-regulated by bacterial effector proteins, it did not provide information on potentially down-regulated miRNAs or which of these miRNAs would be up-regulated in the initial stages of infection, before disease symptoms become apparent. In another study, miR393 was shown to be repressed in response to lipopolysaccharide (LPS) treatment ^26^. LPS is a major component of the outer membrane of Gram-negative bacteria and is a potential inducer of plant defense responses. In this case, a single miRNA was identified in the response, making it unclear whether this response is specific to Gram-negative bacteria infection or a more general stress response.

*Fusarium graminearum* is an ascomycete fungal pathogen, the causative agent of Fusarium head blight (FHB, also known as scab or ear blight) disease in wheat and barley, and ear and stalk rot disease in maize. FHB causes significant yield losses, which can exceed 50% when conditions favor the disease; however, it poses a more significant threat to grain quality and animal and human health ^27^. As the disease progresses, *F. graminearum* produces mycotoxins, such as the trichothecene toxin deoxynivalenol (DON) and the oestrogenic mycotoxin zearalenone (ZEA), which result in reduced grain quality ^28^. Consumption of food and feed produced from toxin-contaminated plants can have serious adverse effects on human and animal health ^29^. Currently, there are no wheat cultivars with resistance to FHB, and limited information is available on the mechanisms employed by the host plant to respond to the disease, as well as how the fungus targets host physiology to promote infection. Therefore, there exists a need to better understand how plants respond to infection by this fungus. Previous studies of genome-wide expression changes in plants infected with *F. graminearum* have focused on changes in protein-coding genes, however small non-coding RNAs were not assessed. These studies identified changes in defense related genes, as well as genes involved in primary metabolism, photosynthesis and transcriptional regulation ^30^. The *Fusarium*-*Arabidopsis* interaction has become an important model pathosystem for characterizing the molecular and physiological basis of plant response to *F. graminearum* ^28–30^. The number of currently annotated miRNA genes in *Arabidopsis thaliana* (approx. 430) is significantly higher than in the cereal species this fungus normally infects ^31^. Therefore, here we investigated changes in plant miRNA expression levels during the early stages (*i.e.*, prior to the onset of visible disease symptoms) of *F. graminearum* infection of *Arabidopsis thaliana* leaves to understand the early plant responses elicited by this fungus.

## Results and Discussion

### Determination of the timing for sampling

As the goal of this work was to detect early changes in miRNA expression in response to *F. graminearum* infection, we first carried out an experiment to determine the time from inoculation to the onset of symptoms. Two 5-week-old Arabidopsis plants were infiltrated with *F. graminearum* and plants were visually inspected twice a day to detect early signs of symptoms. At 5 days, visible chlorosis becomes evident on the inoculated leaves (Supplementary Fig. S1); therefore, we chose 3- and 4-days post inoculation (dpi) as the timepoints to detect early changes in the miRNA transcriptome.

### Differentially expressed miRNAs

RNA-seq analysis of miRNAs was conducted on leaf samples collected from Arabidopsis plants infected with the fungal pathogen *F. graminearum* and mock-inoculated controls at 3-dpi and 4-dpi. We initially conducted an analysis to determine miRNAs that were differentially expressed (DE) when comparing all samples infected with *F. graminearum* with those mock-inoculated (*i.e.*, control). Using this approach, a total of 93 miRNAs (out of 428 precursor miRNAs) were identified to be DE between samples infected with *F. graminearum* and control samples considering 3-dpi and 4-dpi samples together. The complete list of DE miRNAs identified in 3-dpi and 4-dpi vs. control is provided in Supplementary Table S1. Out of these 93 DE miRNAs, 53 miRNAs were up-regulated, and 40 miRNAs were down-regulated (Fig. 1).

**Figure 1.**
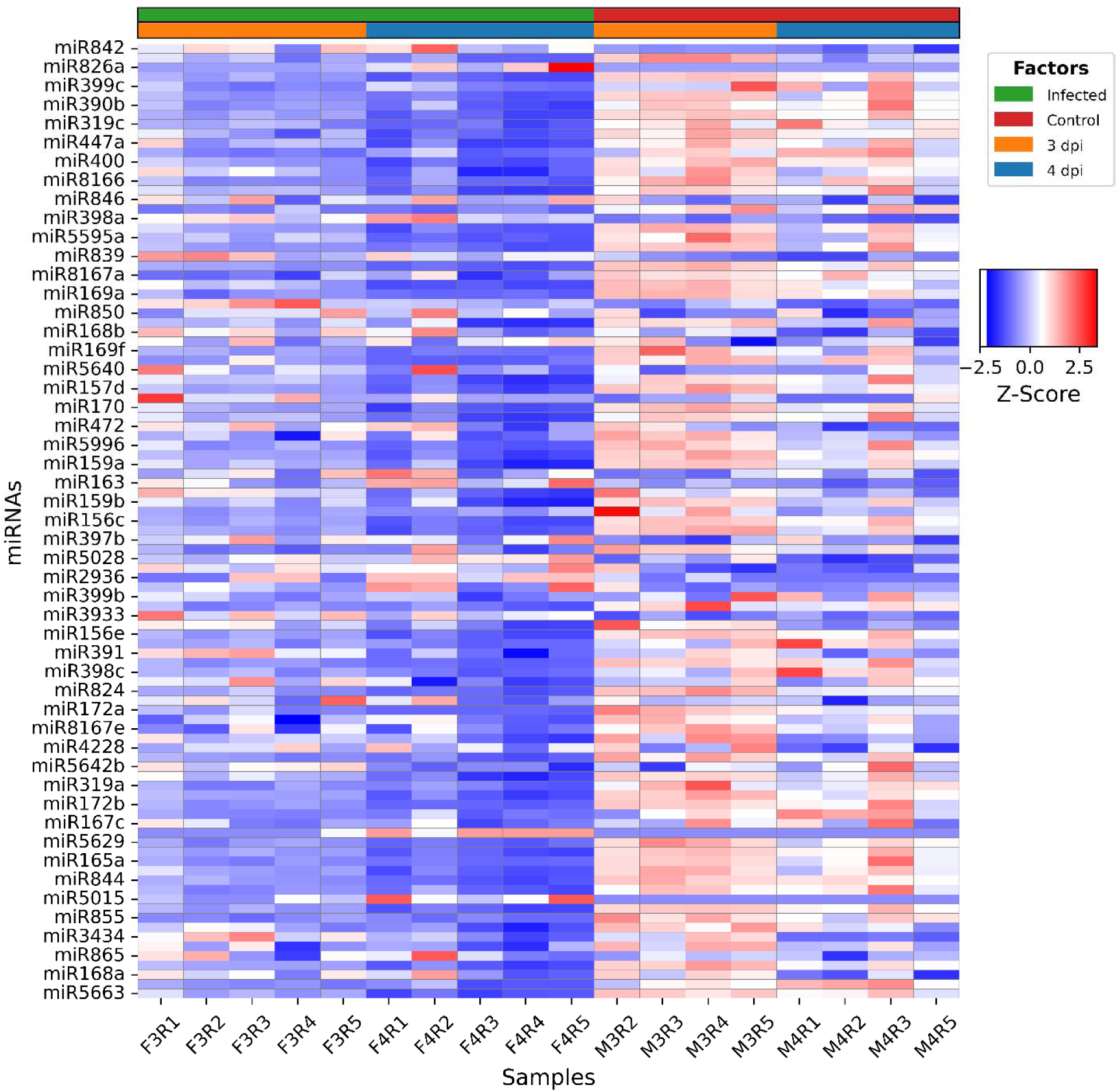
Heatmap of DE miRNAs between infected (3-dpi and 4-dpi combined) and control (3-dpi and 4-dpi combined). Y-axis represents DE miRNAs, and x-axis represents infected and control samples.

To assess miRNA expression changes at 3-dpi and 4-dpi vs. control individually, we also conducted DE analysis for each time point separately. As we did not observe significant differences in the expression of miRNAs in mock-inoculated samples between 3-dpi and 4-dpi, we combined all the control samples into a single set in this analysis (Fig. 2). The complete list of DE miRNAs identified in both 3-dpi infected vs. control and 4-dpi infected vs. control are provided in Supplementary Tables S2 and S3, respectively. In total, there were 65 DE miRNAs when 3-dpi infected samples were compared to the control, whereas 95 miRNAs were DE between 4-dpi infected samples and the control. There were 53 DE miRNAs common to both sampled time points (Fig. 3). Among the 3-dpi fungal infected samples, ten miRNAs (miR842, miR5648, miR3434, miR5651, miR397b, miR5028, miR398a, miR839, miR866, and miR3933) showed at least 2-fold up-regulation relative to the control, whereas only three miRNAs (miR165a, miR855, miR834) showed ≥2-fold down-regulation (Supplementary Table S2). In 4-dpi fungal infected samples, there were 31 DE miRNAs with ≥2-fold up- or down-regulation (6 down-regulated and 25 up-regulated), which included 11 out of the 13 DE miRNAs identified in 3-dpi infected samples (Supplementary Table S3).

**Figure 2.**
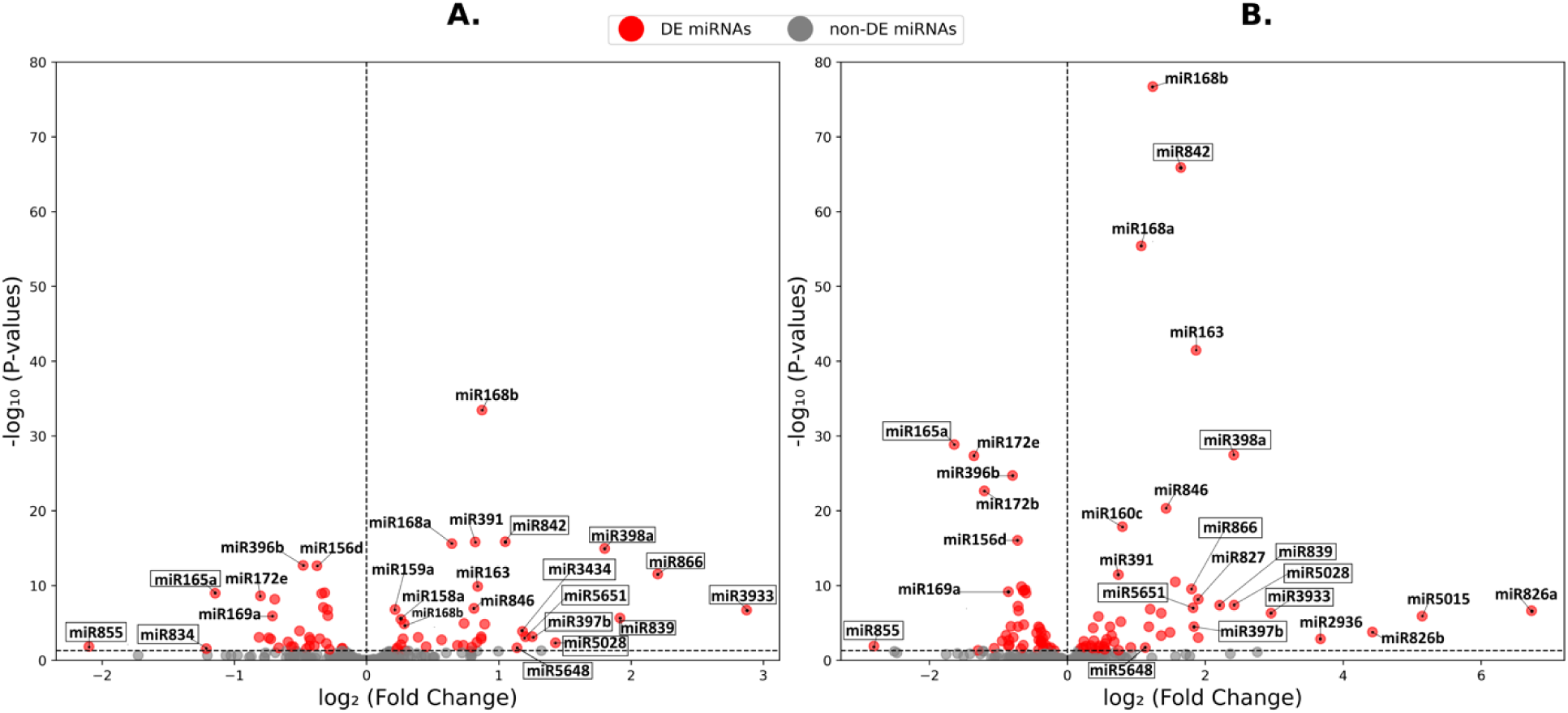
Volcano plot for the DE miRNAs identified in (**A**) 3-dpi infected vs. control, and (**B**) 4-dpi infected vs. control. MiRNAs in rectangular boxes represent common DE miRNA between 3-dpi infected vs. control and 4-dpi infected vs. control with ≥2-fold change.

**Figure 3.**
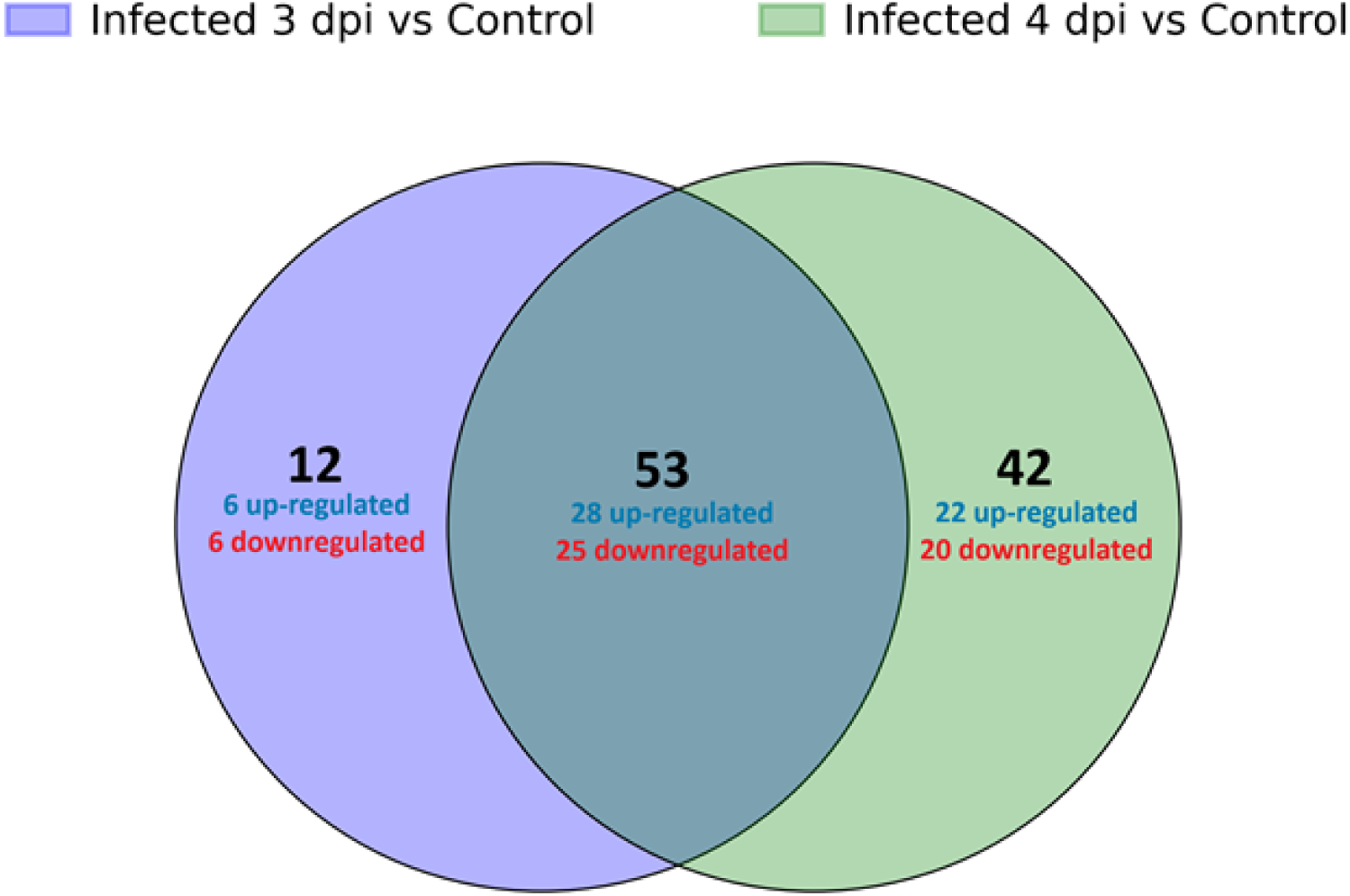
Venn diagram of DE miRNAs identified in 3-dpi infected vs. control and 4-dpi vs. control comparisons.

Of the ten miRNAs that were ≥2-fold up-regulated in 3-dpi infected samples, miR398a has been shown to respond to biotic and abiotic stress and is part of a general stress response mechanism by plants ^32^. MiR397a has been implicated in controlling various plant processes by targeting the mRNAs of genes such as laccases (lignin biosynthesis), β-tubulin, CKB3 (circadian rhythm), and other genes involved in plant growth and development ^33, 34^; miR397a expression has also been shown to respond to copper and cadmium toxicity^35^. Although the up-regulated miRNAs (miR842, miR5648, miR3434, miR5651, miR5028, miR3933, and miR839) have also been identified in other transcriptome analyses, they appeared relatively recently in evolution and their functions have not yet been completely elucidated. Both miR3933 (formerly called miR2328) and miR839 seem to associate with AGO4 (instead of the more common AGO1-associated miRNAs) and can generate small interfering RNAs (siRNAs) that influence DNA methylation of their target genes ^36, 37^.

MiR855 was down-regulated approximately 4.3-fold in 3-dpi and 7-fold in 4-dpi infected samples compared to the control. A miR855 homolog in wheat (Ta-miR855) was identified using a computational approach and predicted to target a MYB transcriptional activator and a transporter in this species in response to cold and salt stresses ^38^. Therefore, it is possible that manipulation of miR855 expression may have direct application in developing wheat cultivars with improved resistance to abiotic and biotic stresses.

MiR826a and miR5015 showed the highest up-regulation in 4-dpi infected plants with 106-fold and 35-fold, respectively (Fig. 2 and Supplementary Table S3). MiR826 is a recently evolved miRNA, which seems to only be present in *A. thaliana*. This miRNA was up-regulated in Arabidopsis plants grown under nutrient deficiency conditions, specifically carbon, nitrogen and sulfur ^39, 40^, whereas it is down-regulated early during exposure of plant roots to toxic levels of copper and cadmium ^41^. Its up-regulation under nitrogen starvation conditions has been linked with a decrease in the synthesis of methionine-derived glucosinolates, which are specialized metabolites also involved in plant defense against biotic stressors ^40^. Therefore, our results showing that miR826a is highly up-regulated in response to fungal infection is surprising and warrants future investigation.

Together with miR156 and miR5021, miR5015 was identified by a computational approach to be involved in trichome development associated with the synthesis of essential oil biosynthesis in mint (*Mentha* spp.) plants ^42^, however no additional information about this miRNA is available. It will be interesting to test the expression of putative targets of miR5015 to identify its role during *F. graminearum* infection of plants.

### Functional analysis of the DE miRNA’s target genes

We performed a Gene Ontology (GO) enrichment analysis to gain an insight over the functional roles of genes that are targeted by the DE miRNAs. We obtained 774 and 724 target genes of DE miRNAs in 3-dpi infected vs. control and 4-dpi infected vs. control from public databases, respectively. We performed GO enrichment analysis of these target genes separately and obtained a total of 189 GO Biological Process (BP), 46 GO Cellular Component (CC), and 80 GO Molecular Function (MF) terms (Table 1), which correspond to the union of two GO enrichment analyses. Over 85% of these GO terms appeared in both enrichment analyses. Enriched GO BP terms included stress-related terms such as defense response to bacterium, regulation of DNA-templated transcription, defense response, leaf senescence, response to abscisic acid, and positive regulation of programmed cell death. The entire list of enriched GO terms is provided in Supplementary Tables S4-S6. Fig. 4 illustrates the top ten BP, CC, and MF enriched GO terms in 3-dpi infected vs. control and 4-dpi infected vs. control comparisons based on GO enrichment False Discovery Rate (FDR).

**Figure 4.**
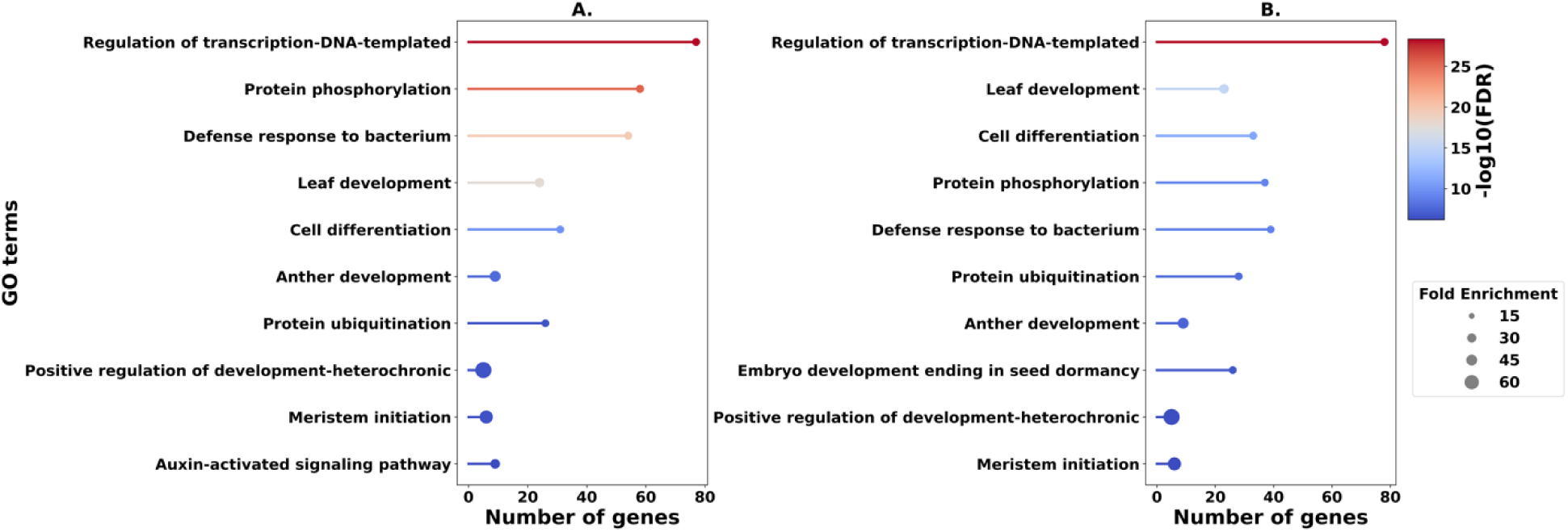
Top 10 enriched GO Biological Process terms for (A) the target genes of DE miRNAs obtained from 3-dpi infected vs. control (B) the target genes of DE miRNAs obtained from 4-dpi infected vs. control.

**Table 1.** Number of enriched GO terms for the target genes of DE miRNAs. The first two columns show the unique number of enriched GO terms for each experiment, followed by the number of common GO terms. *dpi*: days post inoculation.

|  | **3-dpi vs. control** | **4-dpi vs. control** | **Common** |
| --- | --- | --- | --- |
| **Biological Process** | 22 | 17 | 167 |
| **Molecular Function** | 4 | 2 | 69 |
| **Cellular Component** | 6 | 5 | 40 |

Unsurprisingly, plants infected with *F. graminearum* appear to change expression of genes related to transcriptional control and reprogramming development, as well as synthesis of proteins involved in mitigating the effects of oxidative stress. Enrichment for 1,3-β-D-glucan synthase complex likely reflects synthesis of callose (a polymer containing β-1,3-linked glucose) by plants to restrict penetration by the fungus; these polymers are also typical of fungal cell walls ^43^. Callose synthesis and deposition in various tissues has been associated with responses to both biotic and abiotic stresses in plants ^44, 45^.

### Expression changes of miRNA’s target genes

To confirm that the changes in miRNA expression observed in response to *F. graminearum* infection result in the expected modulation of expression of their downstream target genes, we conducted RT-qPCR analyses of a select number of target genes from significantly enriched GO terms (Fig. 5). Gene *AT1G30460* encodes AtCPSF30, a subunit of a cleavage and polyadenylation specificity factor, and is a positive regulator of programmed cell death (PCD) ^46^. This gene is predicted to be targeted by miR157c, which showed 1.4-fold down-regulation in 4-dpi infected samples relative to the control. AtCPSF30 showed similar corresponding up-regulation of 1.6-fold at 3-dpi and 1.8-fold at 4-dpi (Fig. 5A).

**Figure 5.**
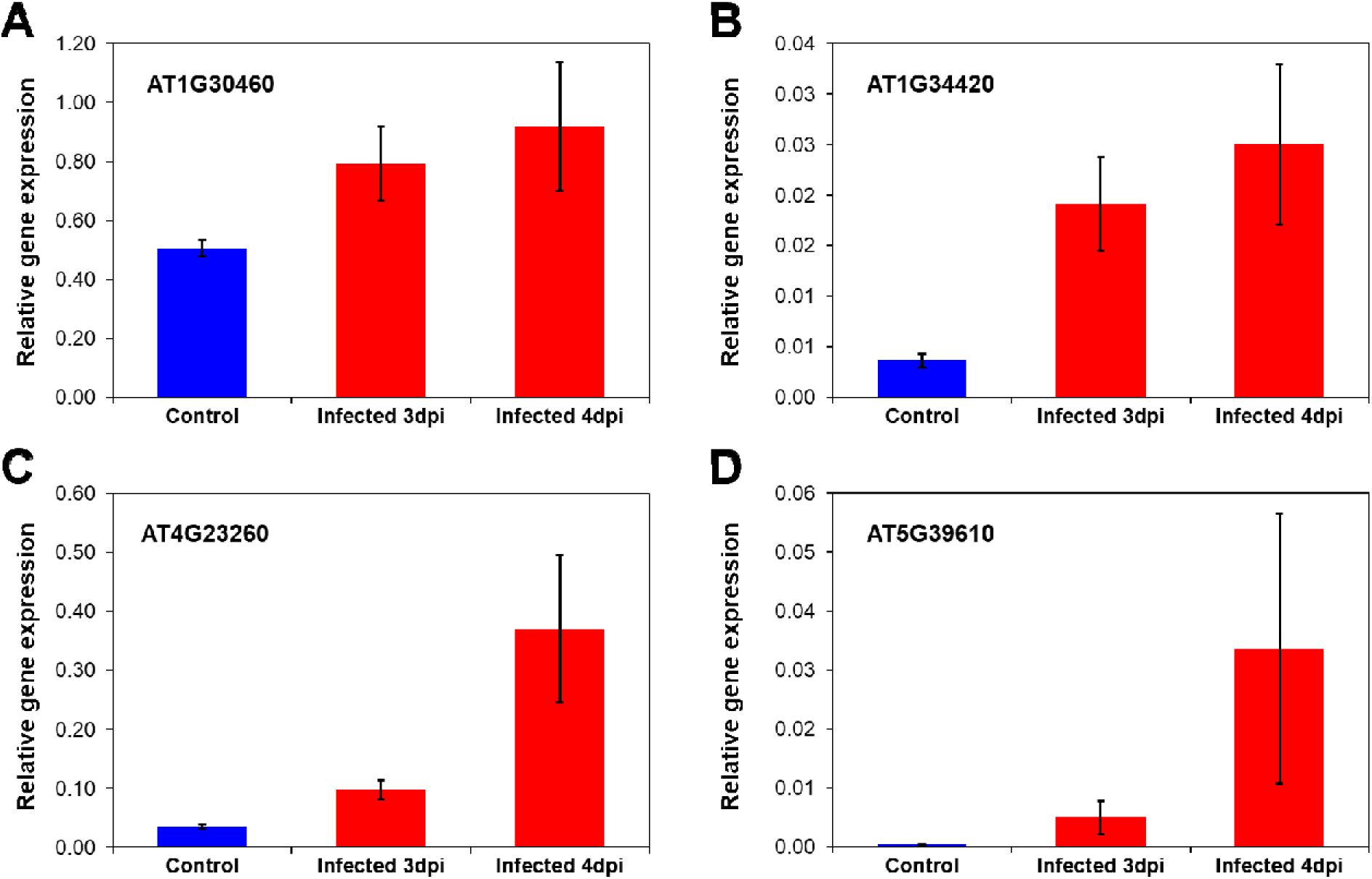
Expression of target genes of top DE miRNAs in 3-dpi and 4-dpi fungal infected, and mock inoculated (control) plants. (A) *AT1G30460*; (B) *AT1G34420*, *BTL2*; (C) *AT4G23260*, *Cysteine-rich receptor-like protein kinase 18*; (D) *AT5G39610, ANAC092.* The expression of PR1 and PR2 was normalized to that of EF1α. The data are shown as mean ± SEM (standard error, n=4).

Gene *AT1G34420* encodes a receptor kinase protein called Bak-to-Life 2 (BTL2), which triggers autoimmunity through activation of the Ca^2+^ channel CNGC20 in a kinase-dependent manner when the PRR (pattern recognition receptor) co-receptors BAK1/SER4 are perturbed ^47^. *BTL2* was up-regulated almost 7-fold in 4-dpi infected samples (Fig. 5B), and is predicted to be targeted by miR396b, which was down-regulated 1.6-fold in 4-dpi samples relative to the control.

Gene *AT4G23260* encodes a cysteine-rich receptor-like protein kinase (AtCRK18) and is a predicted target of miR172c. CRKs play a role in disease resistance and plant cell death ^48^. CRK18 was up-regulated more than 10-fold in 4-dpi infected plants relative to the mock inoculated control plants (Fig. 5C), and miR172c was down-regulated 3.5-fold.

Gene *AT5G39610* encodes a NAC-domain transcription factor (ANAC092), which positively regulates aging-induced cell death and senescence in leaves. This gene has been shown to be upregulated in response to salt stress in Arabidopsis, as well as in response to ABA, ACC and NAA treatment ^49^. ANAC092 is targeted by miR164b, which was down-regulated approximately 1.5-fold in 4-dpi samples. This modest down-regulation of miR164b resulted in a large 91-fold up-regulation of its target gene *ANAC092* (Fig. 5D).

The changes observed in all four miRNA target genes tested were consistent with an up-regulation of the target genes when their corresponding predicted miRNAs are down-regulated in response to fungal infection. Programmed cell death (PCD) is a genetically controlled process that occurs during both plant development and defense responses ^50, 51^. Hypersensitive response is a form of PCD in plants that is linked to resistance to several pathogens, including fungi such as *F. graminearum*. Up-regulation of AtCPSF30, AtCRK18 and ANAC092 is thus consistent with plants inducing PCD to arrest pathogen growth ^52^. BTL2 is a recently identified LRR-RK (leucine-rich repeat receptor kinase) protein that functions as a type of surveillance system that activates multiple signaling pathways to modulate immunity in plants. Increased BTL2 expression has been shown to trigger autoimmunity, which is kept in check by BAK1 phosphorylation of BTL2 ^47^.

Overall, our results indicate that during the early stages in the process of infection by the fungal pathogen *F. graminearum*, Arabidopsis plants respond by changing the levels of miRNAs that control expression of proteins involved in restricting the pathogen’s growth, especially by triggering the hypersensitive response that included programmed cell death. Some of the DE miRNAs identified here, *e.g.*, miRNA855, may inform new approaches to develop plant cultivars with improved resistance to *F. graminearum* infection.

## Methods

### Plant growth

*Arabidopsis thaliana* Col-0 seeds underwent surface sterilization using 10% commercial bleach before being planted on Murashige and Skoog (MS) agar plates. Subsequently, the plates were stored at 4°C for two days and transferred to a growth chamber (Conviron ATC26) with short-day (10-h light/14-h dark) cycles at 150 μmol m^−2^ s^−1^ light intensity, 22°C and 60-70% relative humidity. After 14 days, seedlings were moved to pots filled with Sunshine Mix #1 (Sun Gro Horticulture) and cultivated under identical conditions for an additional 3 weeks.

### Cultivation of *Fusarium graminearum* and plant inoculation

Pathogen cultivation and inoculation was carried out as described in Nalam et al. ^53^. Briefly, *Fusarium graminearum* isolate Z-3639 was grown in ½ strength Potato Dextrose Agar (PDA) plates for 8-10 days at 28°C. Plates were flooded with 10 ml of sterile Milli-Q H_2_O and mycelia was scraped with a plastic cell spreader, carefully avoiding scraping the media. The mycelia suspension was then filtered in four layers of sterile cheesecloth, which was washed with an additional 5 ml of sterile Milli-Q H_2_O. Arabidopsis plants, approximately 5 weeks old, were inoculated with the aid of a 1-ml needleless syringe, by infiltrating the whole leaf surface on the abaxial side, totaling 4 leaves per plant. In parallel, mock plants were infiltrated with Milli-Q H_2_O only. Five plants were inoculated per treatment.

### RNA extraction, library preparation and sequencing

Two infiltrated leaves from each plant (mock or infected) were harvested into a 1.5 ml microcentrifuge tube as a single biological replicate, totaling five replicates per treatment. Samples were flash frozen in liquid N_2_, followed by grinding with a sterile pestle. Total RNA was extracted with TRIzol (ThermoFischer Scientific), following standard protocol. For small RNA library preparation, RNA was quantified with QuBit (ThermoFisher Scientific) Broad Range kit and RNA quality was checked the High Sensitivity RNA ScreenTape Analysis (Agilent). Libraries were generated with the QIAseq miRNA Library Kit (Qiagen) and sequenced using a NextSeq 550 High Output kit at the University of North Texas Health Science Center Genomics Core.

### Processing of smallRNA-seq data

We used FastQC to assess the quality of small RNA-seq data ^54^. We removed adapter sequences using *Trimmomatic* ^55^. On average, 3% of the raw reads were filtered out due to low quality score. Furthermore, we only kept reads of 16–30 nt in length for further analysis to focus on reads that come from miRNAs. We used *bowtie2* ^56^ with *--very-sensitive-local* parameter to map the reads to the *Arabidopsis thaliana* mature miRNA reference transcriptome obtained from mirBase ^31^. To compute read counts for mature miRNAs, we used Samtools ^57^ to process the alignment file and the miRNA annotation file obtained from miRBase. One replicate for each 3-dpi and 4-dpi mock samples were eliminated from the analysis due to low read counts.

### Computing differentially expressed miRNAs

We computed differentially expressed (DE) miRNAs between infected (3-dpi and 4-dpi combined) and control (3-dpi and 4-dpi combined). To observe differential expressions at each time point, we also computed DE miRNAs between 3-dpi infected vs. control and 4-dpi infected vs. control. Because the expression profile did not change between 3 dpi and 4 dpi for the control samples, we combined them for the analyses. For all cases, we used DESeq2 ^58^ with default parameters and reported miRNAs with adjusted p-value ≤ 0.05 as DE. To ensure low expressed miRNAs are not considered for DE analysis, we only kept miRNAs with sum of read counts > 10 across all samples.

### Functional analysis of the DE miRNAs target genes

To investigate the functions of identified DE miRNAs, we performed Gene Ontology (GO) enrichment analysis for the known target genes of DE miRNAs obtained from 3-dpi infected and control and 4-dpi infected vs. control separately. MiRNA target genes were obtained from TarDB ^59^, mirTarBase v9 ^60^ and TarBase v8 ^61^. We utilized AmiGO ^62^ with default parameters and FDR cutoff < 0.05 to obtain significantly enriched GO terms associated with DE miRNA’s target genes.

Quantitative RT-PCR (qRT-PCR) analysis of four miRNA target genes was conducted to confirm the inverse correlation between miRNA levels and expression of their target genes. One microgram of the same total RNA extracted from plants infected with *F. graminearum* or mock inoculated was treated with DNAse and cDNA was synthesized using LunaSCript RT MasterMix kit (New England Biolabs), with random hexamer and oligo-dT primers. Quantitative PCR was carried out in 96-well plates using gene-specific primers and a VWR qPCR Master Mix (Avantor) for 40 cycles on a QuantStudio 6 Flex Real-Time PCR System (ThermoFisher). Relative expression was calculated as described in Hellemans *et al.* ^63^ using the *EF-1* gene (*AT5G60390*) as endogenous reference.

## Acknowledgements

We thank Dr. Taegun Kwon at the Genomics Core (University of North Texas Health Science Center) for library preparation and sequencing. We also thank Dr. Jyoti Shah (University of North Texas) for kindly providing the *Fusarium graminearum* isolate Z-3639 used in this study. This study was supported by a seed grant from the College of Science and College of Engineering at the University of North Texas (to SB and MSA), and the National Institute of General Medical Sciences of the National Institutes of Health under Award Number R35GM133657 (to SB).

## Author contributions

M.S.A. and S.B. conceived the study. S.S.F. and J.H. conducted the experiments, and S.P. analyzed the sequencing data. All authors analyzed the results and prepared the manuscript.

## Data availability statement

All sequencing data from this study are available at the NCBI Sequence Read Archive (SRA) with BioProject accession number PRJNA1136861 and Gene Expression Omnibus (GEO) with accession number GSE272587.

