## Supplementary material for "Early changes in microRNA expression in Arabidopsis plants infected with the fungal pathogen *Fusarium graminearum*": Suppl. Fig. S1

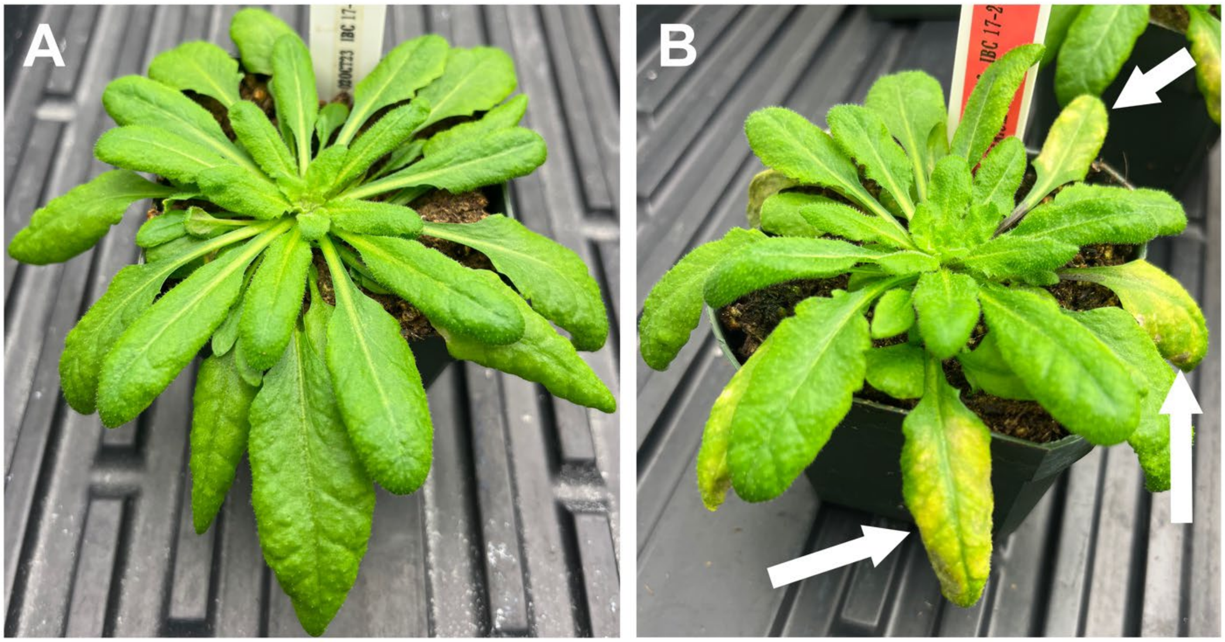


**Supplementary Figure S1:** Preliminary inoculation of Arabidopsis plants with Fusarium graminearum to determine the time for development of symptoms. (**A**) Mock inoculated plant at 5 days post inoculation; (**B**) Fungus inoculated plant at 5 days post inoculation. Arrows point to chlorosis symptom on inoculated leaves.
